# Human germline mutation and the erratic evolutionary clock

**DOI:** 10.1101/058024

**Authors:** Priya Moorjani, Ziyue Gao, Molly Przeworski

## Abstract

Our understanding of the chronology of human evolution relies on the “molecular clock” provided by the steady accumulation of substitutions on an evolutionary lineage. Recent analyses of human pedigrees have called this understanding into question, by revealing unexpectedly low germline mutation rates, which imply that substitutions accrue more slowly than previously believed. Translating mutation rates estimated from pedigrees into substitution rates is not as straightforward as it may seem, however. We dissect the steps involved, emphasizing that dating evolutionary events requires not “a mutation rate,” but a precise characterization of how mutations accumulate in development, in males and females—knowledge that remains elusive.

## Introduction

One of the most fundamental discoveries in evolutionary biology is the “molecular clock”: the observation that changes to the genome along an evolutionary lineage accumulate steadily with time [1-3] and the subsequent development of a theory—the Neutral Theory—that explains why this behavior is expected for neutral genetic changes (i.e., changes with no fitness effects) [4,5]. We now understand that neutral mutations fix in the population at the rate at which they arise, irrespective of demographic history or natural selection at linked sites [4,6]. Thus, the accumulation of neutral substitutions over generations provides a record of the time elapsed on a lineage. It is this “evolutionary clock” that allows researchers to date past events.

Conversely, the existence of an evolutionary clock allows the number of substitutions on a lineage to be translated into a yearly mutation rate, given an independent estimate of when that lineage branched off [2,7-9]. For example, interpreting the fossil record as reflecting a 30 million year (My) split time between humans (apes) and rhesus macaques (Old World Monkeys (OWM)) and using the average nucleotide divergence of ~6.2% between the two species [10] suggests an average yearly mutation rate of 10^-9^ per base pair (bp). Until 2010, single nucleotide substitutions were the main source of data from which to learn about mutation rates, and analyses of substitution patterns consistently suggested rates of around 10^-9^ per bp per year for primates [9,11-13].

Recent findings in human genetics therefore threw a spanner in the works when they suggested de novo mutation rates estimated from human pedigrees to be less than half what was previously believed, or approximately 0.5×10^-9^ per bp per year [14,15]. Because sequencing pedigrees is a much more direct and in principle definitive approach to learn about mutation, these new rate estimates have been widely adopted. They have led to a reappraisal of the chronology of human evolution, suggesting in particular that populations split longer ago than previously believed (e.g., [14,16]). Extrapolating farther back in time becomes problematic however, as pedigree-based estimates imply split times with other primates that are older than compatible with the fossil record, at least as currently interpreted [17-21]. One possible solution, suggested by Scally and Durbin (2012) [14] as well as others, is that yearly mutation rates have decreased towards the present, consistent with the “hominoid rate slowdown” observed in phylogenetic data [22-24].

As we discuss, changes in the yearly mutation rate over the course of human evolution are not only plausible, but follow from first principles. The expected number of de novo mutations inherited by a child depends on paternal (and, to a lesser extent, maternal) ages at puberty and reproduction [25-27], traits that differ markedly among extant primates [18,28,29]. Because these traits evolve, there is no fixed mutation rate per generation, and almost certainly no fixed mutation rate per year. An important implication is that the use of mutations to date evolutionary events requires a precise characterization of how germline mutations accumulate in development, in males and females, and across species. We argue that this knowledge is still elusive and that, as a result, it remains unclear how to set the evolutionary clock. For recent time depths, however, a complementary approach from the study of ancient DNA samples may offer a solution.

In discussing these points, we focus almost exclusively on humans, in part because in other species studied to date, estimates of spontaneous mutations are instead *higher* than substitution rates and the underlying reasons are likely distinct [30-33]. Likewise, we do not discuss mutation rates estimates for mitochondrial DNA; the sources of mutations, complete linkage, and selection pressures make the evolutionary dynamics of this one locus quite distinct from that of the nuclear genome and indeed the discrepancy there too is opposite [34,35]. Moreover, we concentrate on the rate of single nucleotide substitutions in autosomes; for other types of mutations and a discussion of variation in mutation rates along the genome, see [25,36-38].

## The puzzle

Heritable mutations stem from accidental changes to the genome that occur in the development of the germline and production of egg and sperm. A natural definition of the germline mutation rate “per generation” is therefore the rate at which differences arise between the genome of a newly formed zygote and the gametes that it eventually produces. While this quantity cannot be readily measured, it has recently become possible to estimate something highly related, the number of mutations seen in the genome of an offspring’s soma but absent from the parents’ [39] (henceforth *μ_G_*). At least a dozen whole genome studies have applied this approach, resequencing parents and offspring, usually in trios. They reported estimates of *μ_G_* on the order of 10^-8^ per bp (Table 1).

**Table 1.**
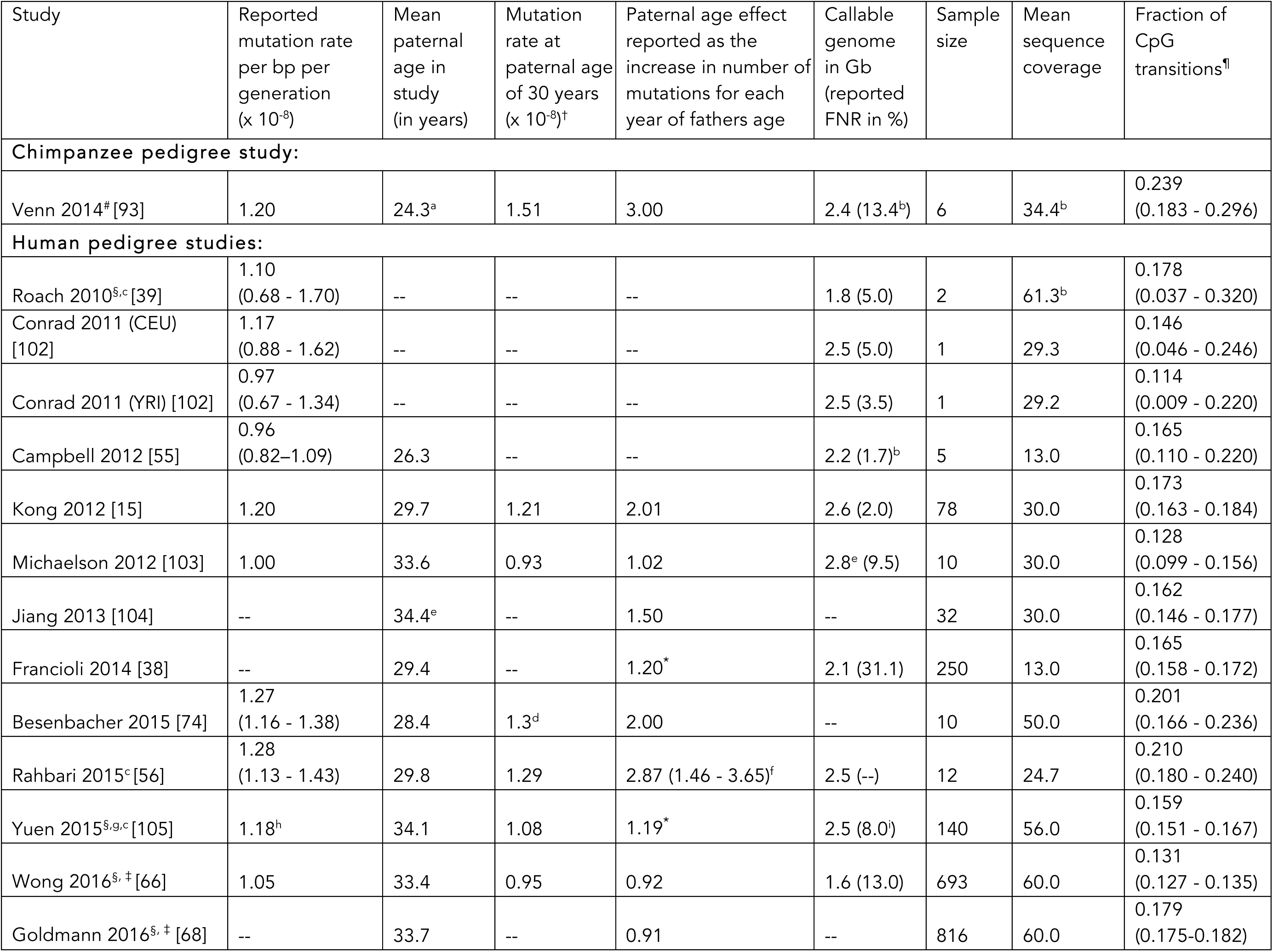
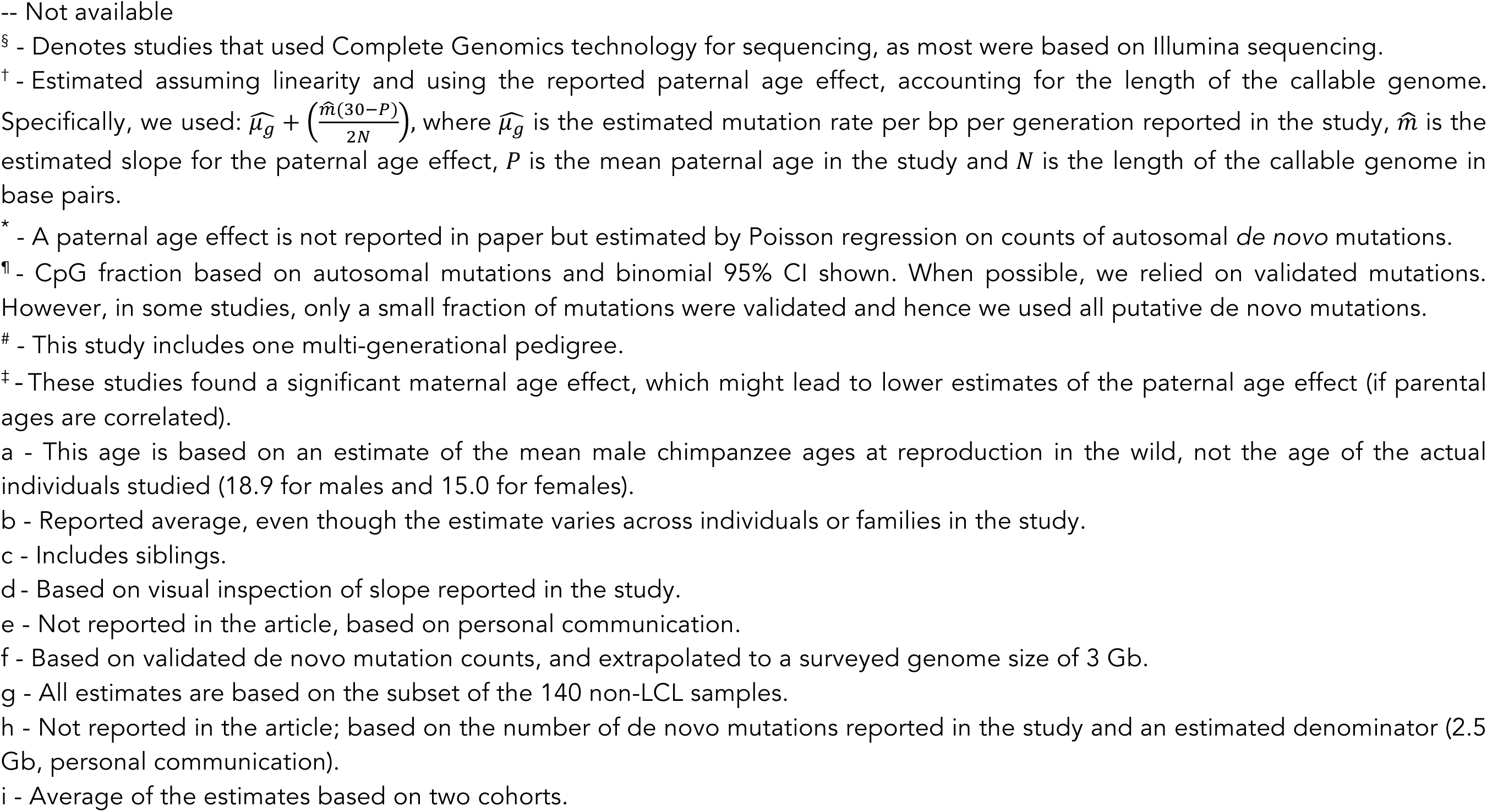
Estimates of mutation rates from pedigree studies

| Study | Reported mutation rate per bp per generation ($\times 10^{-8}$) | Mean paternal age in study (in years) | Mutation rate at paternal age of 30 years ($\times 10^{-8}$)<sup>†</sup> | Paternal age effect reported as the increase in number of mutations for each year of fathers age | Callable genome in Gb (reported FNR in %) | Sample size | Mean sequence coverage | Fraction of CpG transitions<sup>¶</sup> |
| --- | --- | --- | --- | --- | --- | --- | --- | --- |
| **Chimpanzee pedigree study:** |  |  |  |  |  |  |  |  |
| Venn 2014<sup>#</sup> [93] | 1.20 | 24.3<sup>a</sup> | 1.51 | 3.00 | 2.4 (13.4<sup>b</sup>) | 6 | 34.4<sup>b</sup> | 0.239 (0.183 - 0.296) |
| **Human pedigree studies:** |  |  |  |  |  |  |  |  |
| Roach 2010<sup>§,c</sup> [39] | 1.10 (0.68 - 1.70) | -- | -- | -- | 1.8 (5.0) | 2 | 61.3<sup>b</sup> | 0.178 (0.037 - 0.320) |
| Conrad 2011 (CEU) [102] | 1.17 (0.88 - 1.62) | -- | -- | -- | 2.5 (5.0) | 1 | 29.3 | 0.146 (0.046 - 0.246) |
| Conrad 2011 (YRI) [102] | 0.97 (0.67 - 1.34) | -- | -- | -- | 2.5 (3.5) | 1 | 29.2 | 0.114 (0.009 - 0.220) |
| Campbell 2012 [55] | 0.96 (0.82–1.09) | 26.3 | -- | -- | 2.2 (1.7)<sup>b</sup> | 5 | 13.0 | 0.165 (0.110 - 0.220) |
| Kong 2012 [15] | 1.20 | 29.7 | 1.21 | 2.01 | 2.6 (2.0) | 78 | 30.0 | 0.173 (0.163 - 0.184) |
| Michaelson 2012 [103] | 1.00 | 33.6 | 0.93 | 1.02 | 2.8<sup>e</sup> (9.5) | 10 | 30.0 | 0.128 (0.099 - 0.156) |
| Jiang 2013 [104] | -- | 34.4<sup>e</sup> | -- | 1.50 | -- | 32 | 30.0 | 0.162 (0.146 - 0.177) |
| Francioli 2014 [38] | -- | 29.4 | -- | 1.20<sup>*</sup> | 2.1 (31.1) | 250 | 13.0 | 0.165 (0.158 - 0.172) |
| Besenbacher 2015 [74] | 1.27 (1.16 - 1.38) | 28.4 | 1.3<sup>d</sup> | 2.00 | -- | 10 | 50.0 | 0.201 (0.166 - 0.236) |
| Rahbari 2015<sup>c</sup> [56] | 1.28 (1.13 - 1.43) | 29.8 | 1.29 | 2.87 (1.46 - 3.65)<sup>f</sup> | 2.5 (--) | 12 | 24.7 | 0.210 (0.180 - 0.240) |
| Yuen 2015<sup>§,g,c</sup> [105] | 1.18<sup>h</sup> | 34.1 | 1.08 | 1.19<sup>*</sup> | 2.5 (8.0<sup>i</sup>) | 140 | 56.0 | 0.159 (0.151 - 0.167) |
| Wong 2016<sup>§, ‡</sup> [66] | 1.05 | 33.4 | 0.95 | 0.92 | 1.6 (13.0) | 693 | 60.0 | 0.131 (0.127 - 0.135) |
| Goldmann 2016<sup>§, ‡</sup> [68] | -- | 33.7 | -- | 0.91 | -- | 816 | 60.0 | 0.179 (0.175-0.182) |

-- Not available
§ - Denotes studies that used Complete Genomics technology for sequencing, as most were based on Illumina sequencing.
† - Estimated assuming linearity and using the reported paternal age effect, accounting for the length of the callable genome. Specifically, we used: $\widehat{\mu}_g + \left(\frac{\widehat{m}(30-P)}{2N}\right)$ , where $\widehat{\mu}_g$ is the estimated mutation rate per bp per generation reported in the study, $\widehat{m}$ is the estimated slope for the paternal age effect, $P$ is the mean paternal age in the study and $N$ is the length of the callable genome in base pairs.
\* - A paternal age effect is not reported in paper but estimated by Poisson regression on counts of autosomal *de novo* mutations.
¶ - CpG fraction based on autosomal mutations and binomial 95% CI shown. When possible, we relied on validated mutations. However, in some studies, only a small fraction of mutations were validated and hence we used all putative *de novo* mutations.

Although the trio studies were primarily conducted to identify de novo disease mutations, they also inform our understanding of the chronology of primate evolution. Assuming that changes to the genome are neutral, the expected sequence divergence (that is, the expected number of substitutions per bp) between a pair of species, *d*, equals 2*μt*, where *μ* is the mutation rate and *t* is the average time to a common ancestor (i.e., divergence time). Thus, given an estimate of *μ* and orthologous sequences from more than one species, an estimate of *t* can be obtained from *d*/2*μ*. In practice, researchers are interested in an estimate of *t* in years, not generations, and therefore require an estimate of the yearly mutation rate, 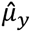. To obtain it, common practice has been to divide the 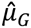 obtained from sequencing of parents and children by a typical age of reproduction (i.e., an estimate of the generation time) [14]. Doing so yields 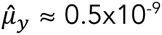 per bp [14,40].

Taken at face value, this mutation rate suggests that African and non-African populations split over 100,000 years [14,16] and a human-chimpanzee divergence time of 12 Mya (for a human-chimpanzee average nucleotide divergence of 1.2% at putatively neutral sites) [10,14,17]. These estimates are older than previously believed, but not necessarily at odds with the existing—and very limited— paleontological evidence for homininae [16,26,41]. More clearly problematic are the divergence times that are obtained for humans and orangutans or humans and OWMs. As an illustration, using whole genome divergence estimates for putatively neutral sites [10] suggests a human-orangutan divergence time of 31 Mya and human-OWM divergence time of 62 Mya. These estimates are implausibly old, implying a human-orangutan divergence well into the Oligocene and OWM-hominoid divergence well into or beyond the Eocene. Thus, the yearly mutation rates obtained from pedigrees seem to suggest dates that are too ancient to be readily reconciled with the current understanding of the fossil record [41,42].

Another way of viewing the same problem is to compare values of 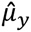 obtained from resequencing pedigrees to those obtained from divergence levels among primates, given estimates of divergence times *t*based on the fossil record. Such estimates of *t* are highly indirect, in part because the fossil record is sparse and in part because relying on fossils with derived traits provides only a lower bound for when the species split [19,20]. A further complication is that for closely related species, *t* reflects the time since the species split as well as the average time to the common ancestor in the ancestral population, which can be substantial [17,43]. Notably, for humans and chimpanzees, the divergence time *t* is thought to be at least 2 million years older than the split time, and possibly much more [17,21,44]. Thus, this approach is mired in uncertainty. Nonetheless, until recently, the consensus in the field has been to use 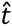 values of 6-7.5 million years ago (Mya) for humans and chimpanzees [9,45], 15-20 Mya for humans and orangutans [46] and 25-35 Mya for humans and OWMs [23,46,47]. Assuming these values and solving 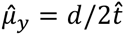 suggests a mutation rate of 10^-9^ per year, more than two-fold higher than what is obtained from pedigree-based estimates. In other words, accepted divergence times suggest that substitutions accumulate faster than they should based on mutation estimates from human pedigrees.

## Could this puzzle be resolved by purifying selection or biased gene conversion?

The equation of mutation and substitution rates is valid only under neutrality. The substitution rate of a population can be factored into two components: the rate at which mutations arise in the population and the probability that a mutation is eventually fixed in the population. When changes are neutral, larger populations experience a greater input of mutations, but exactly counterbalancing this effect is a smaller probability of fixation for each mutation. When natural selection is operating, however, the probability of fixation deviates from the neutral expectation, so the substitution rates at sites under selection are not expected to equal the mutation rate.

To minimize this problem, researchers have focused on putatively neutral regions of the genome when estimating divergence levels (e.g., pseudogenes or genomes with genic and conserved regions excluded) [9,10]. This filtering process is imperfect, however, as putatively neutral regions likely include sites under some degree of natural selection. The net effect should be to decrease the substitution rate relative to the mutation rate, since deleterious mutations that contribute to the count of de novo mutations in pedigrees will not reach fixation (and beneficial alleles are exceedingly rare). In other words, selection should lead to lower substitution rates than mutation rates [34], and likely provides at least a partial explanation for the patterns observed in many taxa (including Drosophila, Arabidopsis and *Caenorhabditis*), as well as for the human mtDNA [31,34]. In contrast, selection only exacerbates the puzzle of why estimated substitution rates are higher than estimated mutation rates in the human nuclear genome.

In addition to selection, the fixation probability can also be affected by GC-biased gene conversion (BGC). This process preferentially resolves mismatches in heteroduplex DNA arising from meiotic recombination in favor of strong alleles (C or G) over weak alleles (A or T), leading to an increased fixation probability of mutations from A/T to C/G and a decreased probability of C/G to A/T mutations relative to neutrality. Although clear evidence for BGC has been observed both in mammalian substitution patterns and in human pedigree data [48,49], the net impact on genome-wide substitution rates remains unclear. Moreover, a similar discrepancy between pedigree and phylogenetic estimates of mutation remains when focusing only on the subset of sites not subject to BGC [10,15]. Thus, this process is unlikely to help reconcile the estimates either.

With no obvious explanation at hand, the surprisingly low mutation rates estimated from pedigrees have led to considerable discussion about whether our understanding of primate evolution is simply incorrect, and divergence times much older than believed. We contend that in some ways this reevaluation is premature. Indeed, while our current understanding of the primate fossil record could be inaccurate, there is underappreciated complexity in the conversion of mutation rates from pedigrees into mutation rates per year and its translation into substitution rates, which remains to be resolved. We try to unpack this complexity, by discussing each step in turn: (1) what it is we are truly estimating from resequencing pedigrees; (2) what we have learned to date and what we have yet to understand; (3) and how to translate the mutation rates into evolutionary dates (Fig 1).

**Fig 1.**
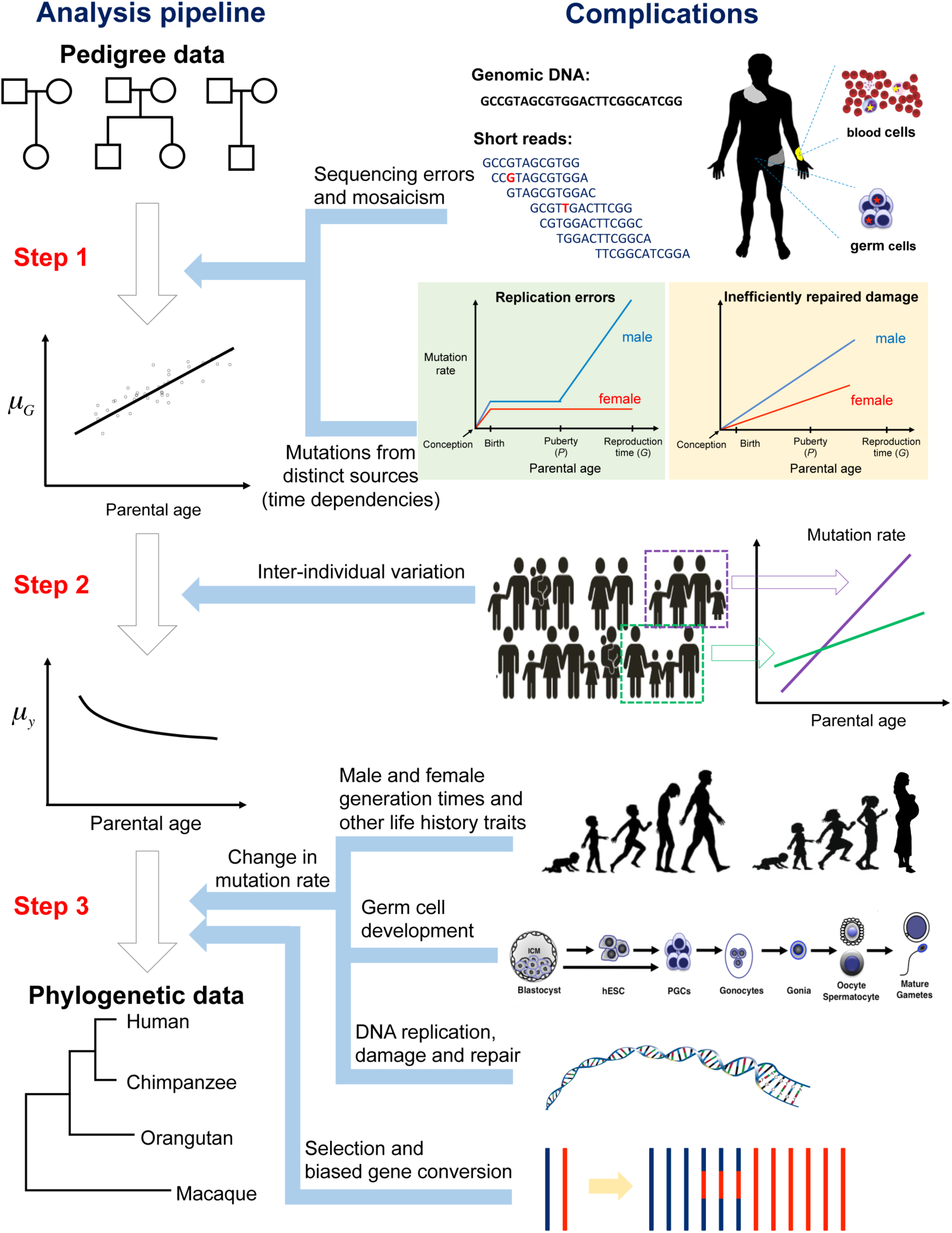
The many steps involved in the conversion of mutation rate estimates from pedigree studies into yearly substitution rates.

## Step1: What exactly is being estimated from human pedigrees?

Human pedigree studies have relied primarily on blood samples from trios, by identifying mutations present in ~50% of reads in the child but absent in both parents. A mutation *rate* is obtained by dividing the count of mutations by the number of base pairs for which there was complete power to identify de novo mutations, or equivalently, dividing it by the genome length, adjusting for power at a typical position in the genome (assuming mutation rates in inaccessible regions of the genome are similar to those in surveyed regions).

Because the mutation rate is so low (~10^-8^ per bp per generation), it is challenging to reliably identify de novo mutations using current sequencing technologies, given the presence of cryptic copy number variation, alignment uncertainty and other confounders [50,51]. Detection pipelines therefore have high false discovery rates, and a stringent set of filters on sequence complexity, read depth, and allelic balance of the reads have to be applied to weed out spurious mutations [52]. This aggressive filtering process substantially increases specificity but decreases the number of sites at which mutations can be detected, so the false negative rate has to be carefully assessed for any given set of filters.

An additional complication is “mosaicism”, that is the presence of two or more genotypes in a given population of cells. When calculating the mutation rate per generation, any mutation accumulated in a germline cycle from zygote to zygote should be included, regardless of the stage at which it occurred (Fig 2, solid stars). The difficulty is that neither the parents nor the offspring are sampled as zygotes; instead, blood samples are used. In these somatic samples, some of the mutations detected will have arisen during the development process of the child, and should not be counted towards germline mutations in the parents (Fig 2 hollow stars; [53]). Moreover, when multiple reads support the alternative allele in the blood of the parent, it is unclear whether the mutation is mosaic and present at high frequency or truly constitutional (i.e., heterozygous in all cells). Standard filtering process requires there to be a balanced number of reads carrying both alleles in the child and no (or very few) reads with the alternative allele in both parents. These criteria will lead to the inclusion of some postzygotic mutations that arose in the child (in germline and/or soma) and the exclusion of a fraction of true germline mutations in the parents (especially those that arose in early development stages) (Fig 2).

**Fig 2.**
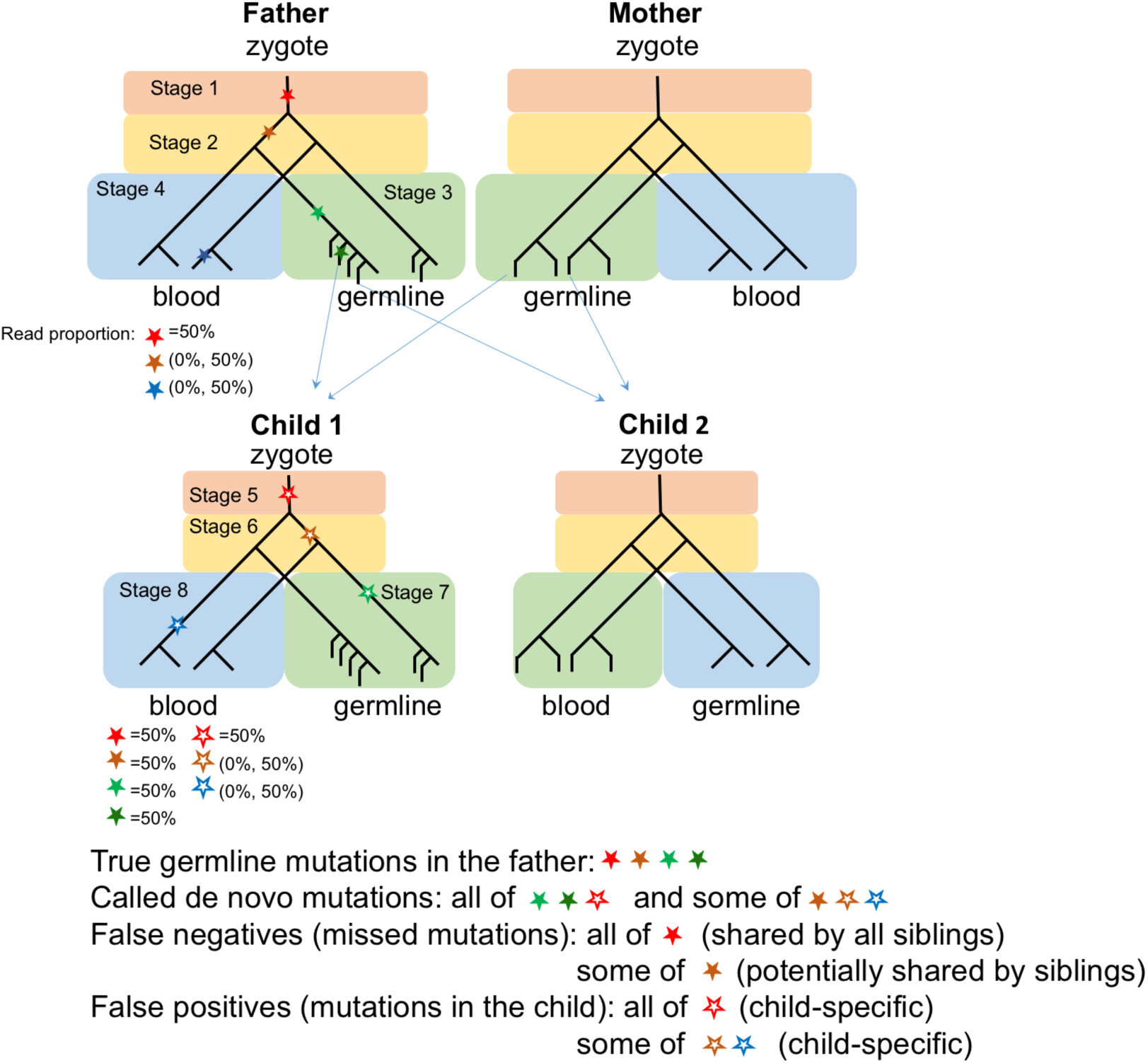
Schematic illustration of mutations occurring during embryonic development and gametogenesis. For simplicity, we show only mutations that arose in the father and one offspring (child 1). Stars represent mutations that originate in different stages of embryogenesis and gametogenesis of the father and the offspring; solid stars are mutations that arise in the father and hollow stars those that occur in the offspring. Shown below each individual are the expected frequencies of the labeled mutations in his or her blood sample. Red, brown and green stars are heritable and should be included in an estimate of germline mutation rates, whereas blue stars are somatic mutations present only in blood samples, which should be excluded. The detection of mutations that are mosaic in both soma and germline strongly suggests that in the cell lineage tree of human development, soma and germline are not reciprocally monophyletic [56,101]. The standard pipelines require allelic balance in the child and no (or very low) read depths in the parents, leading to inclusion of some postzygotic mutations in the child and exclusion of a fraction of germline mutations in the parents. The two effects partially balance, so the overall mutation rate is unlikely to be greatly biased. However, there is a tendency to detect child-specific mutations and to miss ones shared among siblings. As a consequence, the mutation rates during early development are likely underestimated, with potentially important practical implications for predictions of recurrence risk of diseases caused by de novo mutations.

Given the current de novo mutation detection pipelines, the presence of mosaicism therefore leads to two complications: (1) It may lead to a systematic bias in the estimate of the germline mutation rate per generation; and (2) It may distort estimates of per cell division mutation rates in different stages of germline development, due to mis-assignment of the detected mutations to different stages. Current evidence suggests that the first concern is a minor one, both because false negatives and positives are expected to balance out to some extent, and because in practice similar estimates are obtained when considering transmissions in trio studies and when analyzing autozygous segments that descend from a common ancestor multiple generations back (i.e., in which mutations that arose in two or more complete germline cycles are captured) (Table 1; [54,55]). Nonetheless, the current filtering criteria will lead to an underestimate of mosaicism levels and could cloud our understanding of the germline mutational process, impacting the accuracy of predictions about the recurrence risk of diseases caused by de novo mutations [56-58] (see Fig 2).

In addition to these technical considerations, there are conceptual subtleties in interpreting the mutation rate estimates from pedigree studies. As expected a priori and from earlier studies of disease incidences in children [27,59], all large pedigree studies published to date have reported an effect of the age of the father on the total number of de novo mutations inherited by a child (Table 1). Moreover, the increase in the total number of mutations is well approximated by a line [15] (a phenomenon distinct from the few well-studied mutations, such as FGFR2, which occur during spermatogonial stem cell divisions and lead to clonal expansions, and for which the increase in frequency with paternal age is closer to exponential [60-62]). Because spermatogenesis occurs continuously after the onset of puberty, the number of replication-driven mutations inherited by a child is expected to depend on paternal age—more precisely, on the age at which the father enters puberty, his rate of spermatogonial stem cell divisions and age at reproduction [25,63]. Therefore, the observation that the number of mutations increases linearly with paternal age is consistent with a fixed rate of cell division after puberty and a constant rate of mutation per cell division during spermatogenesis.

In contrast, oocytogenesis is completed by the birth of the future mother, so the number of replication-driven mutations inherited by an offspring should be independent of maternal age [64]. For the subset of mutations that do not stem from mistakes during replication—mutations that arise from DNA damage and are poorly repaired for example—there may be a dependence on maternal age as well, if damage accumulates in oocytes [65]. Interestingly, recent studies report that a maternal age effect is also present, potentially supporting the existence of a non-replicative source of germline mutations [66-68]. In any case, what is clear is that the number of de novo mutations in a child is a function of the age of the father at conception and to a lesser extent that of the mother, so values obtained from pedigree studies are estimates of mutation rate *at given mean paternal*(and maternal) ages of the sampled families.

Another complication is that distinct types of mutations may differ in their accrual rates with age, depending on their sources and repair rates over ontogenesis [65,69]. For instance, transitions at methylated CpG sites are thought to occur primarily by spontaneous deamination; beyond this example, the DNA molecule is known to be subject to a large number of chemical assaults from normal cellular metabolism and additional environmental agents [70,71]. While the relative contribution of germline mutations from different sources is unclear, their accrual rates with parental age are unlikely to be identical [68]. Therefore, the mutation rate estimated from pedigree studies is the composite of distinct mutational processes that have distinct dependencies on age and sex [68], making the time-dependency of the overall mutation rate harder to interpret (Fig 1).

With these considerations in mind, what have we learned to date? All large-scale pedigree studies report similar mutation rates per generation, a strong male bias in mutation, and a paternal age effect. On closer inspection, however, their parameter estimates are not consistent. To illustrate this point, we report the estimated mutation rate at paternal age of 30 years, which differ by as much as 40% (Table 1). Given the relatively small sample sizes, some uncertainty is expected from sampling error alone. However, differences in sequencing technology, coverage depth and choice of filters are also likely to be playing a role. As one illustration, the fraction of mutations that involve transitions from CpG sites differs significantly among studies, from 11% to >20% (Chi-Square test, p < 10^-8^, considering studies with a sample size of at least 5). Although biological differences cannot be ruled out, at least some of this variation appears to be due to whether the studies excluded mutations present in dbSNP [72] (because they reasoned that a sequencing error is a more likely explanation than a recurrent mutation). As databases become larger, this step increasingly leads to the exclusion of true mutations [73], with a disproportionate effect on CpG transitions, which are more mutable [74].

Among studies, there is also three-fold variation in the estimated strength of the paternal age effect (Table 1), which remains significant after accounting for the fraction of the genome surveyed for mutation (Fig 3). In principle, differences in the paternal age effect among studies could reflect true biological differences. For instance, a recent study of three larger pedigree families reported that the fathers differed markedly in their paternal age effects (Fig 3) [56]. If indeed fathers differ in the strength of their paternal age effect, then when a single line is fit to data from their offspring, the resulting slope could differ, possibly substantially, from the average slope [25,65]. As the sample size increases, however, the estimated strength of paternal age effect will approach the population mean value, so the observed differences across large studies remain unexplained.

**Fig 3.**
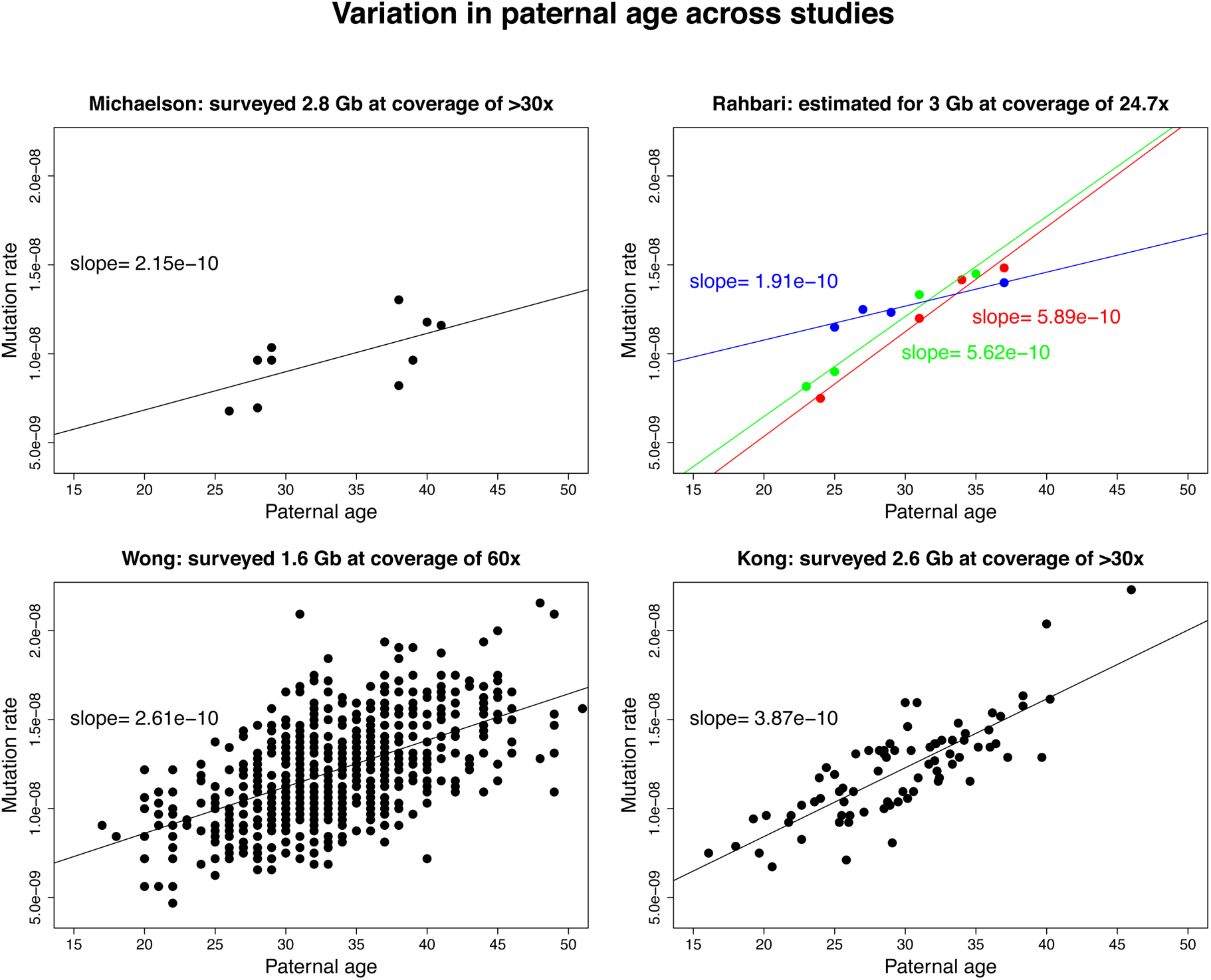
Variation in the estimated paternal age effect, for the autosomes. We plot the de novo mutation rate as a function of the paternal age at conception of the child. The rate was obtained from the reported counts of de novo mutations divided by the fraction of the genome assayed in each study (shown in the title of each subplot, along with the mean coverage per individual). The solid line denotes the fitted slope (i.e., the increase in the mutation rate for each additional year of father’s age). Following the approach of Rahbari et al. 2015 [56], for their study, we used the corrected counts of de novo mutations, which are extrapolated to a genome length of 3 Gb (thereby assuming the mutation rate is the same in the inaccessible region of the genome). The different colors in this study refer to the three different families that they studied: blue, family 244, green, family 603 and red, family 569.

In summary, while pedigree-based approaches are more direct and in principle straight-forward, they have not yet provided a definitive answer about the mutation rate at any given paternal and maternal ages, let alone a precise characterization of how mutations of different sources accumulate over ontogeny in males and females.

## Step 2: How to obtain a yearly de novo mutation rate?

Even if the germline mutation rate per generation, *μ_G_*, were known exactly, strong assumptions would be required to translate the per generation mutation rates of the sampled families into a yearly rate. Common practice has been to obtain a yearly mutation rate by dividing the mutation rate estimated from all the children by the typical age at reproduction (i.e., setting 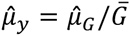, where 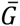 is an estimate of the generation time in the population or the average age of the parents in the study). This practice implicitly ignores differences in reproduction ages across studies, or between the individuals studied and the general population [75]. When such differences exist (Table 1), the values of 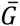 are only comparable if the same 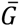 is used and the numerator is replaced by 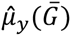, the expected (per generation) mutation rate at a particular age 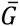. Moreover, if *μ_y_* is not independent of *G*, as suggested by modeling and (limited) available data in humans [25], then it is also important to ask whether the mean parental age for the (predominantly European) samples is representative of the human species, when it is known that ages at reproduction differ substantially across populations [75].

Also complicating matters are possible differences among fathers in the onset of puberty or the strength of their paternal age effect. We know that there is heritable variation among humans (from the same population) in the onset of puberty [76], and there are hints of differences in rate of spermatogonial stem cell divisions or per cell division mutation rates across males [56]. If substantial differences in these factors also exist across human populations, the relationship between *μ_G_* and *G* will vary among them. Therefore, even when the sample size is large and variation in *G* taken into account, the estimated yearly mutation rate from 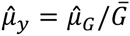 may only be representative of the population(s) under study.

## Step 3: How to relate *μ* to the substitution rate expected over evolutionary time?

### Changes in life history and reproductive traits

Mammalian species vary over three-fold in yearly substitution rates, indicating that the yearly mutation rates change over time [77-79]. In primates, in particular, 35-65% variation is seen in substitution rates across apes and monkeys [10,11]. The cause of variation in substitution rates was long hypothesized to be a “generation time effect”, whereby younger mean ages of reproduction—i.e., shorter generation times—lead to more cell divisions per unit time and hence higher rates of replication-driven mutations [23,80-82]. Support for this claim comes from phylogenetic analyses of mammals, in which reproductive span is the strongest predictor of mutation rates per year among various correlated traits considered, including onset of puberty, body size, metabolic rate, longevity (notably for mitochondrial DNA) and sperm competition [79,83,84].

As we have discussed, a dependence of yearly mutation rates on generation times is expected from what is known about mammalian sperm and egg production. Thus, to accurately convert mutation rates per generation into expected substitution rates per year, changes in the generation time over evolution need to be taken into account. Doing so requires knowledge of numerous parameters that are currently uncertain or simply unknown. A solvable problem is that the conversion depends on the precise dependence of *μ_G_* on parental ages [25], about which there remains considerable uncertainty (Fig 3). A thornier issue is that the yearly substitution rate depends not only on the sex-averaged generation time, but also on the mean ages at reproduction for males and females separately. The reason being that in males, the germline mutation rate depends more strongly on reproductive age than it does in females; thus, for the same average parental age, de novo mutation rates are much lower in a child born to a young father and an old mother than in a child born to an old father and a young mother. As a result, changing the ratio of male to female generation times can have substantial effects on the yearly mutation rate, even when the average remains fixed: for example, a range of ratios from 0.92 to 1.26, as observed in extant hominines, could lead to up to 10% difference in *μ_y_*, and thus introduce uncertainty in phylogenetic dating [26].

Beyond the effect of generation times, the yearly mutation rate will vary with any change in life history traits (e.g., the age at puberty) and germ line developmental process (e.g., the number of cell divisions in each development stage). We know that among extant primates, the onset of puberty differs substantially, from ~1 year in marmosets to 6-13 years in apes [28], as does the length of spermatogonial stem cell divisions [85]. Thus, life history traits can and have evolved across primates. This evolution introduces additional uncertainty in the yearly mutation rate expected at any point in the past [26]. Moreover, these factors influence *μ_y_* in intertwined ways, so it is important to consider their joint effects [10,26].

### Changes in the mutation process

Thus far, we have only discussed sources of changes in the yearly mutation rate due to development and life history, but another layer of evolution occurs at cellular level, in terms of mutational processes of DNA [86-88]. Could the rates of replication error, DNA damage or DNA repair have evolved over millions or even thousands of years? Two recent studies have compared the spectra of rare segregating variants among human populations, and found enrichment of specific mutational signatures in certain populations [88,89]. For instance, Europeans show an increased rate of a TCC -> TTC mutation relative to African or Asian population samples [88]. Additionally, a recent study of autozygous segments in a Pakistani population found some germline mutations to occur at significantly different frequencies than in samples of European ancestry [54]. This observation raises the possibility of recent evolutionary changes in the mutation process itself.

While a change in mutation rates of a specific mutation type is parsimoniously explained by a change in the damage or repair rates, modeling suggests that, even in the absence of such changes, life history traits alone could shift the relative contributions of mutations of different sources [65]. As one example, CpG transitions appear to be more clock-like across species than do other types of mutations (possibly due to a weaker dependence on life history traits) [10,78] and accordingly, the proportion of substitutions due to CpG transitions varies across species [10]. As another example, an increase in paternal age leads not only to an increase in the total germline mutation rate but also to a slight increase in the proportion of mutations in genic regions [38], which should lead to shifts in the mutation spectrum [68]. More generally, it remains highly unclear how much of the differences in mutation rates across populations or species can be attributed to changes in life history and behavior, in the development and renewal of germ cells, in genetic modifiers of mutation (such as enzymes involved in DNA replication and repair) [86,90] or in the environment (such as temperature or the concentration of external mutagens) [91,92].

## Next steps

To predict the rate at which substitutions will accumulate from pedigree data is not as straightforward as it may seem. The main reason being that, while in some contexts a useful concept, there is in fact no such thing as a mutation rate per generation—all that exists is a mean mutation rate for a given set of paternal and maternal life history traits, including ages at puberty and reproduction. These traits are variable among closely related primates [28,29], and heritable variation is seen even among humans [75,76]. Therefore, primate species are expected to differ substantially in both the per generation mutation rate and the yearly mutation rate (e.g., see Table S9 in [26]).

Indeed, phylogenetic analyses show that, over millions of years, substitution rates vary >60% among distantly related primates [10]. The variation in substitution rates across primates and mammals appears to be smaller than that predicted from life history traits in extant species, however [10,26,30]. A likely reason is that, throughout much of their evolutionary past, the lineages had similar life histories. Direct surveys of de novo mutation rates in non-human primates are therefore needed to test whether the present-day mutation rates are more or less similar to those predicted based on life history traits alone.

So far, the only direct estimate of mutation rate in a non-human primate is based on one three-generation pedigree of chimpanzees [93]. The point estimate of the mutation rate at age 30 is higher in chimpanzees than in humans (Table 1), qualitatively consistent with an earlier onset of puberty and faster rate of spermatogenesis [28,85]. Given the differences in detection pipelines, random sampling error and potential intra-species variation, however, these results are still tentative. Both inter- and intra-species variation in mutation rates need to be further characterized in primates.

If mutation rates turn out to vary substantially across species, it will be interesting to examine whether they are well predicted by typical ages at puberty and reproduction. A positive correlation of mutation rate per generation and generation time across species would imply that, over evolutionary timescales, the yearly mutation rate is less variable than the mutation rate per generation, contrary to what is usually assumed (e.g., [81,94]).

If, on the other hand, despite clear differences in life history traits, the per generation mutation rate across mammals turns out to be relatively constant, strong stabilizing selection or developmental constraint must have shaped the evolution of mutation rates. A hint in that direction is provided by recent estimates of mutation rates in mice, whose generation times are on the order of months rather than decades as in apes, and yet whose mutation rate per generation is only about half that of humans [30,95]. It would follow that species with longer generation times will have lower yearly mutation rates, providing direct support for the “generation time effect” than can be obtained from phylogenetic evidence [23,77,81].

That yearly mutation rates are expected to be unsteady poses difficulties for the use of substitutions to date evolutionary events. One solution is to explicitly model the changes in life-history traits over the course of primate evolution and to study their impact on substitution rates. To this end, Amster and Sella (2016) [26] proposed a model that estimates divergence and split times across species, accounting for differences in sex-specific life history and reproductive traits. A next step will be to extend their model to consider replicative and non-replicative mutations separately. In addition, as more reliable estimates of mutational parameters become available from pedigree studies of humans and non-human primates, models will need to be revised to account for differences in cell division rates and possible differences in repair rates. Unfortunately, however, some uncertainty will remain due to lack of knowledge about life history traits in ancestral lineages.

An alternative might be to use only CpG transitions for dating. This solution is based on the observation that CpG transitions accumulate in a quasi-clocklike manner across primates [10,78], as well as across human populations [88]. Puzzlingly, however, in human pedigree data, there is no detectable difference between the effects of paternal age on CpG transitions and on other types of mutations [15,56], suggesting that CpG transitions are no more clock-like. In that regard, it will be highly relevant to compare accrual rates of CpG transitions in large pedigree studies from multiple primate species.

In addition to the use of pedigree studies, two other types of approaches have been introduced recently to learn about mutation rates. The first is a set of ingenious methods that use population genetic modeling to estimate mutation rates based on segments of the genome inherited from a distant common ancestor [96,97]. Unfortunately, these methods rely on a number of other parameter estimates, including a demographic model (on which times to the common ancestor are based), fine-scale meiotic recombination rates or, to obtain yearly rates, generation times. In a related approach, mutation rates are estimated from autozygous segments that descend from a recent common ancestor that can be more reliably inferred. The approach presents a number of advantages, notably in minimizing the possible contribution of somatic mutations, but only provides a mutation rate averaged over both sexes and several generations [54,55].

It has also become possible recently to use reliably-dated ancient DNA samples to estimate average yearly mutation rates over different evolutionary periods. Here, the divergence from an extant sample (e.g., human) to an outgroup (e.g., chimpanzee) is compared to what is seen between an ancient genome and the outgroup. The “missing sequence divergence” then provides an estimate of the average mutation rate per year over that timescale. Applied to archaic human samples from the past 50,000 years, this approach suggests yearly rates around 0.5×10^-9^ per bp [98]. As in the “tip calibration” approach for estimating the evolutionary rates using sequentially sampled virus genomes or ancient mitochondrial genomes [99], the study of many such ancient nuclear samples distributed across ancestral populations could in principle serve as “spike ins” for the evolutionary clock, allowing one to adjust for changes in rates over different time periods along a lineage.

Together, this combination of approaches will both inform us about how to reliably set the evolutionary clock and provide a first direct look at the evolution of mutation rates.

## Acknowledgments

We are grateful to Kay Prüfer and David Reich for organizing “The Human Mutation Rate Meeting” at the Max Planck Institute for Evolutionary Anthropology in February 2015, and to all the participants for many enlightening discussions. We also thank Augustine Kong, Laurent Francioli, Paul de Bakker, Raheleh Rahbari, Ryan Yuen and Wendy Wong for providing data used in generating Table 1 and Fig 3, David Pilbeam for helpful discussions about the fossil record and Michel Georges for helpful discussions about the impact of mosaicism. We are grateful to Nick Barton, Alexey Kondrashov and two anonymous reviewers for their comments on an earlier version of the manuscript, as well as to Guy Amster, Laurent Francioli, Kelley Harris, Iain Mathieson, David Pilbeam, Jonathan Pritchard, and Guy Sella. This study makes use of data generated by the Genome of the Netherlands Project [100]. Funding for the project was provided by the Netherlands Organization for Scientific Research under award number 184021007, dated July 9, 2009 and made available as a Rainbow Project of the Biobanking and Biomolecular Research Infrastructure Netherlands (BBMRI-NL). Samples where contributed by LifeLines (http://lifelines.nl/lifelines-research/general), The Leiden Longevity Study (http://www.healthy-ageing.nl; http://www.langleven.net), The Netherlands Twin Registry (NTR: http://www.tweelingenregister.org), The Rotterdam studies, (http://www.erasmus-epidemiology.nl/rotterdamstudy) and the Genetic Research in Isolated Populations program (http://www.epib.nl/research/geneticepi/research.html#gip). The sequencing was carried out in collaboration with the Beijing Institute for Genomics (BGI).

## References

1. Zuckerkandl E, Pauling L (1962) Molecular disease, evolution, and genic heterogeneity. In: Kasha M, Pullman B, editors. Horizons in Biochemistry Academic Press, New York,. pp. 189–225.

2. Kumar S (2005) Molecular clocks: four decades of evolution. Nat Rev Genet 6: 654–662.

3. Bromham L, Penny D (2003) The modern molecular clock. Nat Rev Genet 4: 216–224.

4. Kimura M (1983) The neutral theory of molecular evolution.: Cambridge University Press.

5. Jukes TH, Cantor CR (1969) Evolution of protein molecules. In: Munro HN, editor. Mammalian protein metabolism New York: Academic Press. pp. pp. 21–123.

6. Birky CW, Jr, Walsh JB (1988) Effects of linkage on rates of molecular evolution. Proc Natl Acad Sci U S A 85: 6414–6418.

7. Kumar S, Subramanian S (2002) Mutation rates in mammalian genomes. Proceedings of the National Academy of Sciences 99: 803–808.

8. Kondrashov FA, Kondrashov AS (2010) Measurements of spontaneous rates of mutations in the recent past and the near future. Philosophical Transactions of the Royal Society of London B: Biological Sciences 365: 1169–1176.

9. Nachman MW, Crowell SL (2000) Estimate of the mutation rate per nucleotide in humans. Genetics 156: 297–304.

10. Moorjani P, Amorim CEG, Arndt PF, Przeworski M (2016) Variation in the molecular clock of primates. bioRxiv: 036434.

11. Yi S, Ellsworth DL, Li W-H (2002) Slow molecular clocks in Old World monkeys, apes, and humans. Molecular Biology and Evolution 19: 2191–2198.

12. Steiper ME, Young NM (2006) Primate molecular divergence dates. Mol Phylogenet Evol 41: 384–394.

13. (2005) Initial sequence of the chimpanzee genome and comparison with the human genome. Nature 437: 69–87.

14. Scally A, Durbin R (2012) Revising the human mutation rate: implications for understanding human evolution. Nat Rev Genet.

15. Kong A, Frigge ML, Masson G, Besenbacher S, Sulem P, et al. (2012) Rate of de novo mutations and the importance of father’s age to disease risk. Nature 488: 471–475.

16. Schiffels S, Durbin R (2014) Inferring human population size and separation history from multiple genome sequences. Nature genetics 46: 919–925.

17. Prado-Martinez J, Sudmant PH, Kidd JM, Li H, Kelley JL, et al. (2013) Great ape genetic diversity and population history. Nature 499: 471–475.

18. Langergraber KE, Prüfer K, Rowney C, Boesch C, Crockford C, et al. (2012) Generation times in wild chimpanzees and gorillas suggest earlier divergence times in great ape and human evolution. Proceedings of the National Academy of Sciences 109: 15716–15721.

19. Steiper ME, Young NM (2008) Timing primate evolution: lessons from the discordance between molecular and paleontological estimates. Evolutionary Anthropology: Issues, News, and Reviews 17: 179–188.

20. Jensen-Seaman MI, Hooper-Boyd KA (2013) Molecular Clocks: Determining the Age of the Human–Chimpanzee Divergence. eLS.

21. Scally A, Dutheil JY, Hillier LW, Jordan GE, Goodhead I, et al. (2012) Insights into hominid evolution from the gorilla genome sequence. Nature 483: 169–175.

22. Goodman M (1962) Evolution of the immunologic species specificity of human serum proteins. Human Biology 34: 104–150.

23. Li W-H, Tanimura M (1987) The molecular clock runs more slowly in man than in apes and monkeys. Nature 326: 93–96.

24. Goodman M (1961) The role of immunochemical differences in the phyletic development of human behavior. Human Biology 33: 131–162.

25. Ségurel L, Wyman M, Przeworski M (2014) Determinants of mutation rate variation in the human germline. Annual Review of Human Genetics 15: 47–70.

26. Amster G, Sella G (2016) Life history effects on the molecular clock of autosomes and sex chromosomes. Proc Natl Acad Sci U S A 113: 1588–1593.

27. Crow JF (2006) Age and sex effects on human mutation rates: an old problem with new complexities. J Radiat Res 47 Suppl B: B75–82.

28. Dixson AF (2009) Sexual selection and the origins of human mating systems: OUP Oxford.

29. Gage TB (1998) The comparative demography of primates: with some comments on the evolution of life histories. Annual Review of Anthropology: 197–221.

30. Uchimura A, Higuchi M, Minakuchi Y, Ohno M, Toyoda A, et al. (2015) Germline mutation rates and the long-term phenotypic effects of mutation accumulation in wild-type laboratory mice and mutator mice. Genome research 25: 1125–1134.

31. Exposito-Alonso M, Becker C, Schuenemann VJ, Reitter E, Setzer C, et al. (2016) The rate and effect of de novo mutations in natural populations of Arabidopsis thaliana. bioRxiv: 050203.

32. Lynch M (2010) Evolution of the mutation rate. Trends Genet 26: 345–352.

33. Smeds L, Qvarnstrom A, Ellegren H (2016) Direct estimate of the rate of germline mutation in a bird. Genome Research: gr. 204669.204116.

34. Ho SY, Lanfear R, Bromham L, Phillips MJ, Soubrier J, et al. (2011) Time-dependent rates of molecular evolution. Molecular ecology 20: 3087–3101.

35. Pulquerio MJ, Nichols RA (2007) Dates from the molecular clock: how wrong can we be? Trends in Ecology & Evolution 22: 180–184.

36. Campbell CD, Eichler EE (2013) Properties and rates of germline mutations in humans. Trends Genet 29: 575–584.

37. Shendure J, Akey JM (2015) The origins, determinants, and consequences of human mutations. Science 349: 1478–1483.

38. Francioli LC, Polak PP, Koren A, Menelaou A, Chun S, et al. (2015) Genome-wide patterns and properties of de novo mutations in humans. Nature genetics 47: 822–826.

39. Roach JC, Glusman G, Smit AF, Huff CD, Hubley R, et al. (2010) Analysis of genetic inheritance in a family quartet by whole-genome sequencing. Science 328: 636–639.

40. Scally A (2016) The mutation rate in human evolution and demographic inference. bioRxiv: 061226.

41. Begun DR (2015) Fossil Record of Miocene Hominoids. Handbook of Paleoanthropology: Springer. pp. 1261–1332.

42. Hartwig WC (2002) The primate fossil record: Cambridge University Press.

43. Mailund T, Munch K, Schierup MH (2014) Lineage sorting in apes. Annual review of genetics 48: 519–535.

44. Wall JD (2003) Estimating ancestral population sizes and divergence times. Genetics 163: 395–404.

45. Patterson N, Richter DJ, Gnerre S, Lander ES, Reich D (2006) Genetic evidence for complex speciation of humans and chimpanzees. Nature 441: 1103–1108.

46. Perelman P, Johnson WE, Roos C, Seuánez HN, Horvath JE, et al. (2011) A molecular phylogeny of living primates. PLoS Genet 7: e1001342.

47. Steiper ME, Young NM, Sukarna TY (2004) Genomic data support the hominoid slowdown and an Early Oligocene estimate for the hominoid–cercopithecoid divergence. Proceedings of the National Academy of Sciences of the United States of America 101: 17021–17026.

48. Duret L, Galtier N (2009) Biased gene conversion and the evolution of mammalian genomic landscapes. Annual review of genomics and human genetics 10: 285–311.

49. Williams AL, Genovese G, Dyer T, Altemose N, Truax K, et al. (2015) Non-crossover gene conversions show strong GC bias and unexpected clustering in humans. Elife 4: e04637.

50. Li H (2014) Toward better understanding of artifacts in variant calling from high-coverage samples. Bioinformatics 30: 2843–2851.

51. Nielsen R, Paul JS, Albrechtsen A, Song YS (2011) Genotype and SNP calling from next-generation sequencing data. Nature Reviews Genetics 12: 443–451.

52. Beal MA, Glenn TC, Somers CM (2012) Whole genome sequencing for quantifying germline mutation frequency in humans and model species: Cautious optimism. Mutation Research/Reviews in Mutation Research 750: 96–106.

53. Scally A (2015) Mutation rates and the evolution of germline structure. bioRxiv: 034298.

54. Narasimhan VM, Rahbari R, Scally A, Wuster A, Mason D, et al. (2016) A direct multi-generational estimate of the human mutation rate from autozygous segments seen in thousands of parentally related individuals. bioRxiv: 059436.

55. Campbell CD, Chong JX, Malig M, Ko A, Dumont BL, et al. (2012) Estimating the human mutation rate using autozygosity in a founder population. Nature genetics 44: 1277–1281.

56. Rahbari R, Wuster A, Lindsay SJ, Hardwick RJ, Alexandrov LB, et al. (2015) Timing, rates and spectra of human germline mutation. Nature Genetics.

57. Acuna-Hidalgo R, Bo T, Kwint MP, van de Vorst M, Pinelli M, et al. (2015) Post-zygotic point mutations are an underrecognized source of de novo genomic variation. The American Journal of Human Genetics 97: 67–74.

58. Behjati S, Huch M, van Boxtel R, Karthaus W, Wedge DC, et al. (2014) Genome sequencing of normal cells reveals developmental lineages and mutational processes. Nature 513: 422–425.

59. Haldane JB (1935) The rate of spontaneous mutation of a human gene. Journal of Genetics 31: 317–326.

60. Maher G, Goriely A, Wilkie A (2014) Cellular evidence for selfish spermatogonial selection in aged human testes. Andrology 2: 304–314.

61. Risch N, Reich E, Wishnick M, McCarthy J (1987) Spontaneous mutation and parental age in humans. American journal of human genetics 41: 218.

62. Arnheim N, Calabrese P (2016) Germline Stem Cell Competition, Mutation Hot Spots, Genetic Disorders, and Older Fathers. Annual review of genomics and human genetics.

63. Crow JF (2000) The origins, patterns and implications of human spontaneous mutation. Nat Rev Genet 1: 40–47.

64. Drost JB, Lee WR (1995) Biological basis of germline mutation: comparisons of spontaneous germline mutation rates among drosophila, mouse, and human. Environmental and molecular mutagenesis 25: 48–64.

65. Gao Z, Wyman MJ, Sella G, Przeworski M (2016) Interpreting the Dependence of Mutation Rates on Age and Time. PLoS Biol 14: e1002355.

66. Wong WS, Solomon BD, Bodian DL, Kothiyal P, Eley G, et al. (2016) New observations on maternal age effect on germline de novo mutations. Nature communications 7.

67. McRae JF, Clayton S, Fitzgerald TW, Kaplanis J, Prigmore E, et al. (2016) Prevalence, phenotype and architecture of developmental disorders caused by de novo mutation. bioRxiv: 049056.

68. Goldmann JM, Wong WS, Pinelli M, Farrah T, Bodian D, et al. (2016) Parent-of-origin-specific signatures of de novo mutations. Nature Genetics.

69. Alexandrov LB, Stratton MR (2014) Mutational signatures: the patterns of somatic mutations hidden in cancer genomes. Curr Opin Genet Dev 24: 52–60.

70. Lodish H (2008) Molecular cell biology: Macmillan.

71. Lindahl T, Wood RD (1999) Quality control by DNA repair. Science 286: 1897–1905.

72. Sherry ST, Ward M-H, Kholodov M, Baker J, Phan L, et al. (2001) dbSNP: the NCBI database of genetic variation. Nucleic acids research 29: 308–311.

73. Lek M, Karczewski K, Minikel E, Samocha K, Banks E, et al. (2015) Analysis of protein-coding genetic variation in 60,706 humans. bioRxiv: 030338.

74. Besenbacher S, Liu S, Izarzugaza JM, Grove J, Belling K, et al. (2015) Novel variation and de novo mutation rates in population-wide de novo assembled Danish trios. Nature communications 6.

75. Fenner JN (2005) Cross-cultural estimation of the human generation interval for use in genetics-based population divergence studies. American journal of physical anthropology 128: 415–423.

76. Day FR, Helgason H, Chasman DI, Rose LM, Loh P-R, et al. (2016) Physical and neurobehavioral determinants of reproductive onset and success. Nature genetics.

77. Wu C-I, Li W-H (1985) Evidence for higher rates of nucleotide substitution in rodents than in man. Proceedings of the National Academy of Sciences 82: 1741–1745.

78. Hwang DG, Green P (2004) Bayesian Markov chain Monte Carlo sequence analysis reveals varying neutral substitution patterns in mammalian evolution. Proc Natl Acad Sci U S A 101: 13994–14001.

79. Wilson Sayres MA, Venditti C, Pagel M, Makova KD (2011) Do variations in substitution rates and male mutation bias correlate with life-history traits? A study of 32 mammalian genomes. Evolution 65: 2800–2815.

80. Wu CI, Li WH (1985) Evidence for higher rates of nucleotide substitution in rodents than in man. Proc Natl Acad Sci U S A 82: 1741–1745.

81. Laird CD, McCONAUGHY BL, McCARTHY BJ (1969) Rate of fixation of nucleotide substitutions in evolution.

82. Kohne DE, Chiscon J, Hoyer B (1972) Evolution of primate DNA sequences. Journal of Human Evolution 1: 627–644.

83. Nabholz B, Glémin S, Galtier N (2008) Strong variations of mitochondrial mutation rate across mammals—the longevity hypothesis. Molecular biology and evolution 25: 120–130.

84. Gillooly JF, Allen AP, West GB, Brown JH (2005) The rate of DNA evolution: effects of body size and temperature on the molecular clock. Proceedings of the National Academy of Sciences of the United States of America 102: 140–145.

85. Ramm SA, Stockley P (2010) Sperm competition and sperm length influence the rate of mammalian spermatogenesis. Biol Lett 6: 219–221.

86. Britten RJ (1986) Rates of DNA sequence evolution differ between taxonomic groups. Science 231: 1393–1398.

87. Thomas GW, Hahn MW (2014) The human mutation rate is increasing, even as it slows. Molecular biology and evolution 31: 253–257.

88. Harris K (2015) Evidence for recent, population-specific evolution of the human mutation rate. Proceedings of the National Academy of Sciences 112: 3439–3444.

89. Mathieson I, Reich DE (2016) Variation in mutation rates among human populations. bioRxiv: 063578.

90. Sharp NP, Agrawal AF (2012) Evidence for elevated mutation rates in low-quality genotypes. Proceedings of the National Academy of Sciences 109: 6142–6146.

91. Lindgren D (1972) The temperature influence on the spontaneous mutation rate. Hereditas 70: 165–177.

92. Lupu A, Pechkovskaya A, Rashkovetsky E, Nevo E, Korol A (2004) DNA repair efficiency and thermotolerance in Drosophila melanogaster from ‘Evolution Canyon’. Mutagenesis 19: 383–390.

93. Venn O, Turner I, Mathieson I, de Groot N, Bontrop R, et al. (2014) Strong male bias drives germline mutation in chimpanzees. Science 344: 1272–1275.

94. Lehtonen J, Lanfear R (2014) Generation time, life history and the substitution rate of neutral mutations. Biology letters 10: 20140801.

95. Adewoye AB, Lindsay SJ, Dubrova YE, Hurles ME (2015) The genome-wide effects of ionizing radiation on mutation induction in the mammalian germline. Nature communications 6.

96. Lipson M, Loh P-R, Sankararaman S, Patterson N, Berger B, et al. (2015) Calibrating the human mutation rate via ancestral recombination density in diploid genomes. PLoS Genet 11: e1005550.

97. Palamara PF, Francioli LC, Wilton PR, Genovese G, Gusev A, et al. (2015) Leveraging Distant Relatedness to Quantify Human Mutation and Gene-Conversion Rates. The American Journal of Human Genetics 97: 775–789.

98. Fu Q, Li H, Moorjani P, Jay F, Slepchenko SM, et al. (2014) Genome sequence of a 45,000-year-old modern human from western Siberia. Nature 514: 445–449.

99. Rieux A, Balloux F (2016) Inferences from tip-calibrated phylogenies: a review and a practical guide. Molecular ecology 25: 1911–1924.

100. Genome of the Netherlands Consortium. (2014) Whole-genome sequence variation, population structure and demographic history of the Dutch population. Nature Genetics 46: 818–825.

101. Bunyan DJ, Robinson DO, Collins AL, Cockwell AE, Bullman HM, et al. (1994) Germline and somatic mosaicism in a female carrier of Duchenne muscular dystrophy. Human genetics 93: 541–544.

102. Conrad DF, Keebler JE, DePristo MA, Lindsay SJ, Zhang Y, et al. (2011) Variation in genome-wide mutation rates within and between human families. Nat Genet 43: 712–714.

103. Michaelson JJ, Shi Y, Gujral M, Zheng H, Malhotra D, et al. (2012) Whole-genome sequencing in autism identifies hot spots for de novo germline mutation. Cell 151: 1431–1442.

104. Jiang Y-h, Yuen RK, Jin X, Wang M, Chen N, et al. (2013) Detection of clinically relevant genetic variants in autism spectrum disorder by whole-genome sequencing. The American Journal of Human Genetics 93: 249–263.

105. Yuen RK, Thiruvahindrapuram B, Merico D, Walker S, Tammimies K, et al. (2015) Whole-genome sequencing of quartet families with autism spectrum disorder. Nature medicine 21: 185–191.

